# Phenotype projections accelerate biobank-scale GWAS

**DOI:** 10.1101/2023.11.20.567948

**Authors:** Michael Zietz, Undina Gisladottir, Kathleen LaRow Brown, Nicholas P. Tatonetti

## Abstract

Understanding the genetic basis of complex disease is a critical research goal due to the immense, worldwide burden of these diseases. Pan-biobank genome-wide association studies (GWAS) provide a powerful resource in complex disease genetics, generating shareable summary statistics on thousands of phenotypes. Biobank-scale GWAS have two notable limitations: they are resource-intensive to compute and do not inform about hand-crafted phenotype definitions, which are often more relevant to study. Here we present Indirect GWAS, a summary-statistic-based method that addresses these limitations. Indirect GWAS computes GWAS statistics for any phenotype defined as a linear combination of other phenotypes. Our method can reduce runtime by an order of magnitude for large pan-biobank GWAS, and it enables ultra-rapid (roughly one minute) GWAS on hand-crafted phenotype definitions using only summary statistics. Overall, this method advances complex disease research by facilitating more accessible and cost-effective genetic studies using large observational data.

## 1 Introduction

Performing GWAS across large-scale resources like the UK Biobank can be challenging and time-consuming. Hurdles to this research include restricted access to personally-identifying genetic and health information, resource-intensive computations, necessary expertise in defining phenotype data and filtering genetic data, and evolving GWAS bestpractices. Pan-biobank GWAS—studies in which GWAS are run on hundreds to thousands of phenotypes—provide a solution to these challenges [1, 2, 3, 4]. These studies pre-compute and publicly release GWAS summary statistics for most or all phenotypes in a dataset, eliminating the need for many GWAS to be duplicated. Pan-biobank GWAS provide a useful starting point for genetic analysis of complex phenotypes, but they have two major limitations: they are highly resource intensive to compute [5, 6], and they only release summary statistics on a limited number of predefined phenotypes—typically single data fields. As a consequence, any new GWAS method, phenotype definition, or cohort selection requires GWAS to be re-computed, representing a major cost to research.

We developed Indirect GWAS, a method that aims to resolve these two limitations. Indirect GWAS is a mathematical method for computing GWAS summary statistics for a phenotype defined as a linear combination of other phenotypes. Because our method is restricted to linear GWAS, this can be done using only summary information. This approach has two aims: to increase the speed of pan-biobank GWAS and to enable researchers to compute GWAS for custom phenotype definitions using only pan-biobank GWAS summary statistics. Indirect GWAS enables large speed improvements for pan-biobank GWAS, potentially reducing computation time by an order of magnitude for the largest pan-biobank studies by reducing the dimension of phenotype data. Additionally, Indirect GWAS enables researchers to approximate GWAS for arbitrarily-defined phenotypes using pan-biobank GWAS summary statistics at very low computational cost. Together, these improvements accelerate the computation of pan-biobank GWAS and increase the usefulness of pan-biobank GWAS results for studying specific traits and diseases.

## 2 Results

### 2.1 Indirect GWAS

Consider a genetic association test for a single phenotype (*y*) and a single genetic variant (*g*), *y ∼ g*. Since this is univariate least squares, assuming both have zero mean, the coefficient solution and standard error are

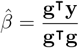

and

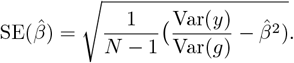

Write the phenotype *y* as a linear combination of *m* feature phenotypes *x*_1_, …, *x*_*m*_, (i.e. **y** = **Xp**). Define two *m*-vectors **b** and **s**, that hold the coefficient and standard error estimates for regressions of the genetic variant against the feature phenotypes (e.g. *b*_*i*_ = **g**^*T*^**x**_*i*_*/***g**^*T*^**g**). Rearranging each standard error term, we can write the Var(*g*) for each feature association test as

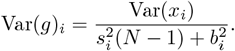

Denote the mean across features as Var(*g*) = Var(*g*)_*i*_ and the covariance matrix of the feature phenotypes as **C**. Then the coefficient and standard error for *y* can be written in terms of feature summary statistics as

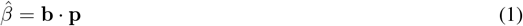

and

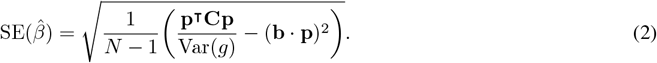

A more comprehensive derivation including covariates is included in the supplementary materials.

### 2.2 Validation in real data

We demonstrated the mathematical correctness of Indirect GWAS by comparing direct to indirect regressions for several GWAS methods using data from the UK Biobank (described in the supplementary methods). First, we randomly sampled 100,000 individuals, 500,000 variants, and 10 quantitative phenotypes, and we generated 10 random projections of these phenotypes. Then, we computed GWAS summary statistics for the 10 random phenotypes using both direct and indirect approaches, applying four different GWAS methods: univariate OLS (only genotype and intercept), multivariate OLS (genotype, intercept, age, sex, 10 genetic PCs), FastGWA [7] (with covariates), and Regenie [8] (with covariates).

For each method, we compared GWAS chi-squared statistics to verify that Indirect GWAS produces equivalent results. Indirect GWAS produced accurate summary statistics for all GWAS methods (*R*^2^ *>* 0.97, Figure 1), indicating that Indirect GWAS is mathematically correct and applicable to both linear and linear mixed models. The small errors in FastGWA and Regenie summary statistics are due to the fact that these methods are only approximately OLS.

**Figure 1.**
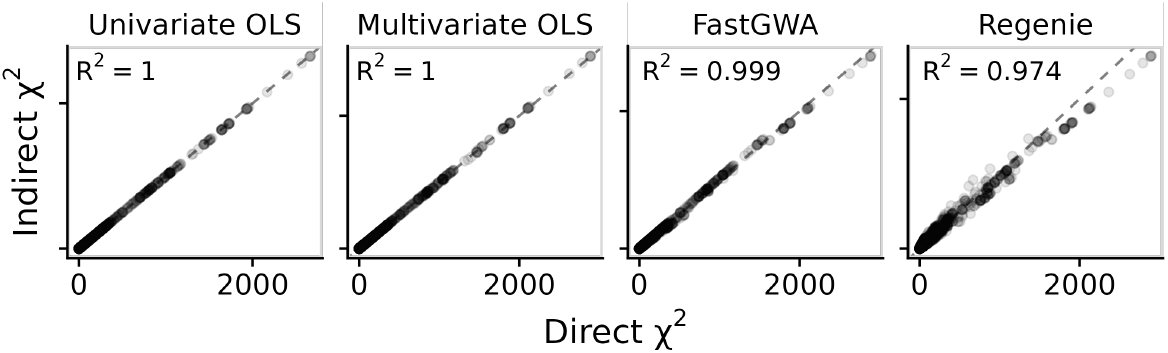
Validation of Indirect GWAS using four GWAS methods. GWAS chi-squared statistics are compared between direct and indirect approaches for 10 randomly projected phenotypes, 500,000 genetic variants using data from the UK Biobank. Each point represents one genetic variant and one phenotype (5,000,000 total points in each facet). *R*^2^ values indicate goodness of fit between the direct and indirect statistics.

### 2.3 Increased pan-biobank GWAS speed

Indirect GWAS affords an approximation to traditional pan-biobank GWAS when phenotypes are correlated (Figure 3A). To do so, we reduce the phenotype dimensionality (e.g. using PCA), perform direct GWAS on the latent space, then reconstruct full-dimensional summary statistics using Indirect GWAS. By reducing the number of GWAS that must be computed using slower methods like LMM, our approach can save a large fraction of computation time.

**Figure 2.**
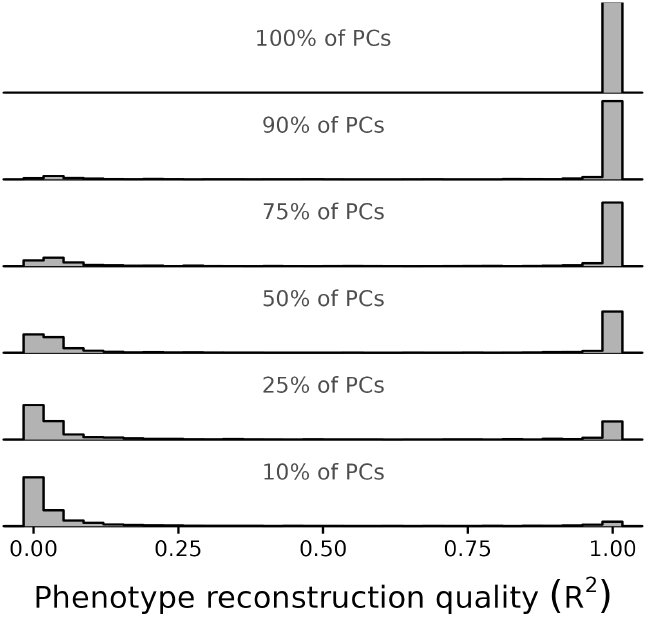
Real phenotypes can be compressed using a small fraction of PCs. 1238 real binary phenotypes from the UK Biobank are shown (ICD-10 codes). As more PCs are retained, the fraction of phenotypes that are well-reconstructed increases.

**Figure 3.**
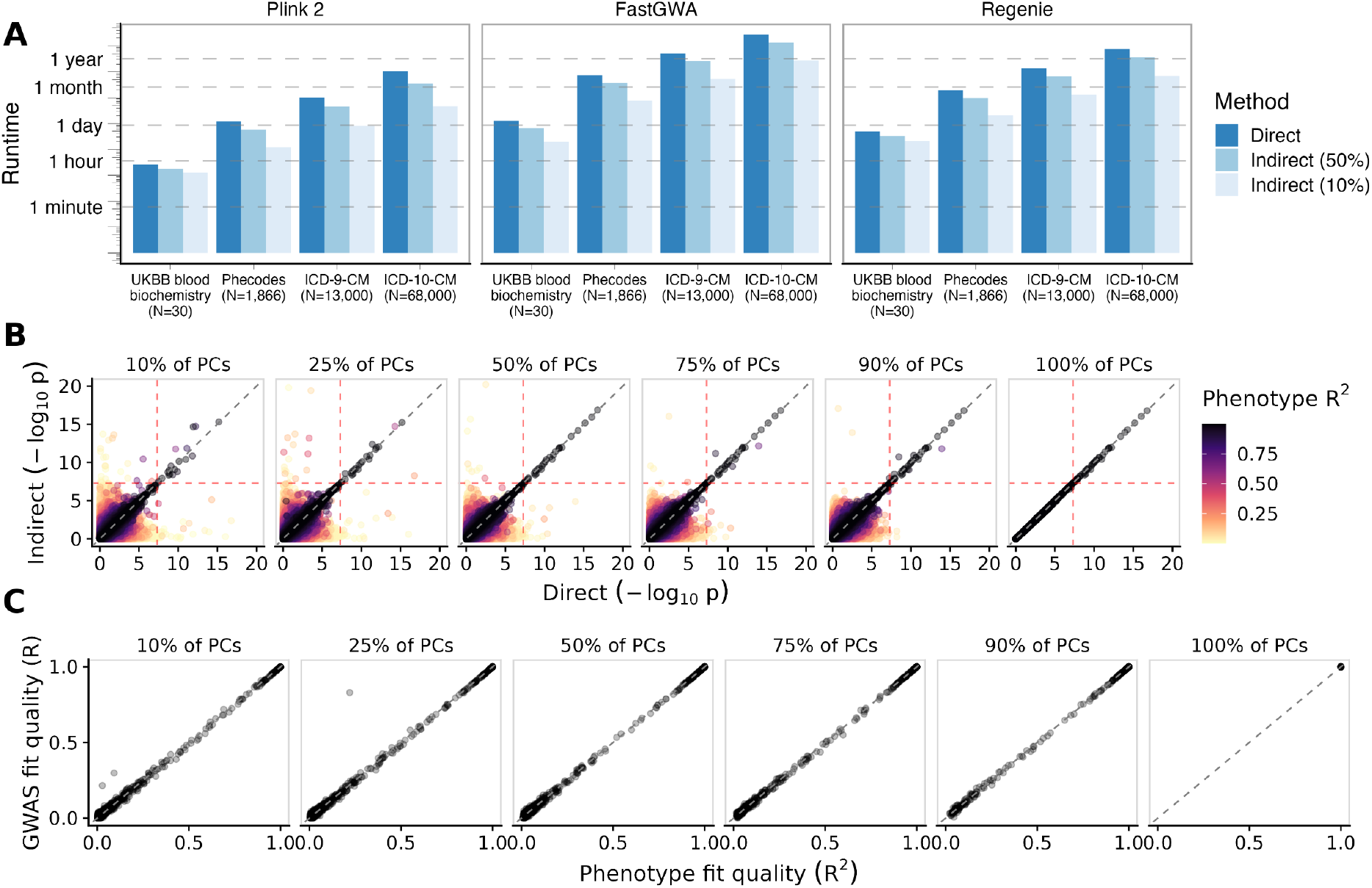
Indirect GWAS accelerates pan-biobank GWAS. **A**. Indirect GWAS reduces computation time for GWAS across many phenotypes. Runtimes are quoted for a cohort of 100,000 individuals and 500,000 genetic variants. Time savings are proportional to the reduction in phenotype dimension. **B**. Indirect GWAS summary statistics better approximate direct summary statistics as more PCs are used. Red lines indicate the traditional cutoff for genome-wide statistical significance, 5 *×* 10^*−*8^. Each point represents one phenotype-variant association. **C**. Phenotype fit predicts Indirect GWAS performance. Phenotype fit is the *R*^2^ value between the phenotype and its reconstruction from a reduced set of PCs. GWAS performance is the Pearson correlation between direct and indirect GWAS negative log p-values. Each point represents one phenotype.

In order to achieve a good GWAS approximation using a low-dimensional latent space, it is important that the phenotypes themselves be well-reconstructed from low dimensions. We evaluated how well 1238 binary phenotypes (ICD-10 codes) from UK Biobank could be compressed using PCA. We found that a large fraction (roughly 50%) of these phenotypes could be reconstructed with *R*^2^ *>* 0.8 using 50% of the PCs, and roughly 73% with *R*^2^ *>* 0.9 using 5% of the PCs (Figure 2).

A larger latent dimension leads to better reconstruction of both phenotypes through PCA and GWAS summary statistics through Indirect GWAS (Figure 3B). We found that Indirect GWAS is able to approximate statistically significant variant p-values using even very small fractions of the PCs, achieving very high recall (*>* 0.98) at all PC dimensions and precision that improves as the latent dimension increases (Table 1).

**Table 1:**
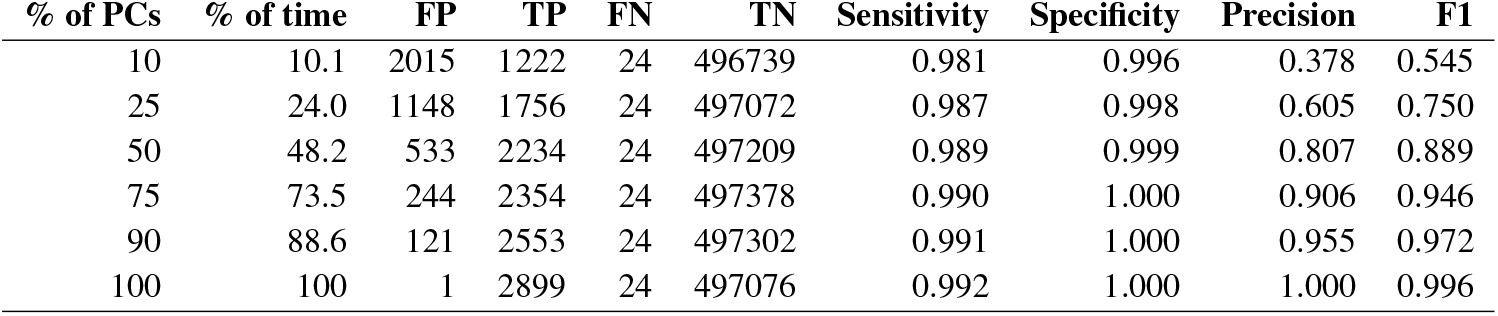
Indirect GWAS performance across various dimensionality reductions. This analysis considered 500,000 variants, 342,350 samples, 1238 ICD-10 phenotypes, and used Plink 2 for GWAS. Variants were considered genome-wide significant if their p-values were less than 5*×*10^*−*8^. “% of time” indicates what fraction of the full, direct runtime would be needed for the indirect approach. FP, TP, FN, and TN indicate false positive, true positive, false negative, and true negative.

| % of PCs | % of time | FP | TP | FN | TN | Sensitivity | Specificity | Precision | F1 |
| --- | --- | --- | --- | --- | --- | --- | --- | --- | --- |
| 10 | 10.1 | 2015 | 1222 | 24 | 496739 | 0.981 | 0.996 | 0.378 | 0.545 |
| 25 | 24.0 | 1148 | 1756 | 24 | 497072 | 0.987 | 0.998 | 0.605 | 0.750 |
| 50 | 48.2 | 533 | 2234 | 24 | 497209 | 0.989 | 0.999 | 0.807 | 0.889 |
| 75 | 73.5 | 244 | 2354 | 24 | 497378 | 0.990 | 1.000 | 0.906 | 0.946 |
| 90 | 88.6 | 121 | 2553 | 24 | 497302 | 0.991 | 1.000 | 0.955 | 0.972 |
| 100 | 100 | 1 | 2899 | 24 | 497076 | 0.992 | 1.000 | 1.000 | 0.996 |

We found that Indirect GWAS can substantially reduce computation time for pan-biobank GWAS (Figure 3A). The time costs for PCA and Indirect GWAS are small compared to the runtime savings from computing fewer direct GWAS. In fact, the computation time savings are roughly in line with the latent space fraction used (Table 1). Overall, these results show that Indirect GWAS can accelerate pan-biobank GWAS in cases where some degree of approximation is acceptable, and the time savings can be immense. Tables S1 and S2 give more detailed runtime results for direct and indirect methods, respectively. Additional details about the runtime measurements are provided in the supplementary methods.

### 2.4 Summary statistics for arbitrary phenotypes

The final goal of this work is to unlock pan-biobank GWAS summary statistics for the analysis of custom phenotype definitions. To address this challenge, we propose approximating a linear map from the biobank phenotypes to the custom phenotype definition using linear regression, reconstructing the summary statistics with Indirect GWAS, and using the regression goodness-of-fit to predict the fidelity of the resulting GWAS summary statistics.

To illustrate this method, we considered phecodes, which group ICD codes into higher-level categories [9]. To explore the effects of more complex phenotype definitions, we compared phecodes using only inclusion criteria to those using both inclusion and exclusion criteria. Phecode inclusion criteria are a simple maximum operation across ICD-10 codes (e.g. phecode 499: cystic fibrosis is present if a patient has any ICD-10 codes that start with E84). Exclusion criteria are designed to exclude individuals from the control group when they are not cases but have codes that are very similar to case inclusion codes and could represent mislabeling (e.g. exclude type 1 diabetes mellitus patients from type 2 diabetes mellitus control group). Inclusion/exclusion definitions are equivalent to inclusion definitions among individuals who are included in the analysis, but they result in different cohorts for each phenotype, thereby violating the linearity assumption of Indirect GWAS in a plausible scenario akin to real genetic analyses.

We used linear regression to approximate a map from the 2616 binary and quantitative phenotypes to 567 phecodes (for inclusion and inclusion/exclusion separately). For each phecode, we evaluated the phenotypic goodness-of-fit using *R*^2^, then computed GWAS summary statistics using both direct and indirect approaches, each run with both inclusion and inclusion/exclusion definitions. Overall, both types of phecodes were well-fit using the features, with all phenotype fit *R*^2^ values over 0.8. The missingness introduced by exclusion criteria reduced the correspondence of the GWAS summary statistics slightly (*R*^2^ value of the negative log p-values reduced from 0.96 to 0.94 by adding exclusion codes, Figure 4). Overall, though, both were well-fit, indicating that Indirect GWAS can approximate GWAS for new phenotypes using only summary statistics.

**Figure 4.**
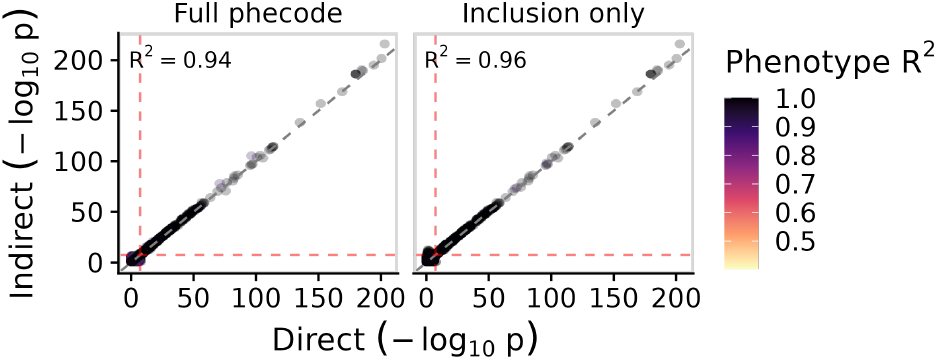
Indirect GWAS performance for phecodes. Shown are GWAS summary statistics for 567 phecodes generated using both direct GWAS and Indirect GWAS using 2616 binary and quantitative phenotypes as features. *R*^2^ values indicate goodness of fit between the direct and indirect summary statistics. Full phecodes use both inclusion and exclusion codes.

### 2.5 Sensitivity to poorly-fit phenotypes

Hand-crafted phenotype definitions used in observational research are often nonlinear, violating the Indirect GWAS model. Nonetheless, many phenotypes can be approximated sufficiently using a linear model that Indirect GWAS may be appropriate. For example, while none of the phecodes are strictly linear functions of ICD codes, their definitions can be approximated by linear models sufficiently well that Indirect GWAS summary statistics match well (Figure 4).

We evaluated how robust Indirect GWAS is to poorly-fit phenotype definitions by comparing phenotype fit quality (measured using *R*^2^) to GWAS summary statistic correspondence (measured using the Pearson correlation between negative log p-values of direct and indirect GWAS methods). Figure 3C shows that phenotype fit quality is an excellent predictor of the quality of Indirect GWAS summary statistics. Phenotypes that can be fit well using linear models will likely have high-quality Indirect GWAS summary statistics.

### 2.6 Performance across categories

We wanted to understand whether Indirect GWAS performs well only for certain phenotypes. Because such a large number of performance measurements are possible, and because we sought to evaluate the influence of the compression fraction as well, we first plotted the performance of Indirect GWAS for every phenotype as a function of the compression fraction (Figure 5A). In this comparison, the performance of Indirect GWAS is computed as the *R*^2^ value between negative log transformed p-values. This is a reasonable metric since Indirect GWAS seeks to reconstruct the direct summary statistics as faithfully as possible. We found that phenotypes uniformly improved their GWAS performance as the compression fraction grew. Still, there were clear differences between phenotypes in what compression fraction was needed to produce good results.

**Figure 5.**
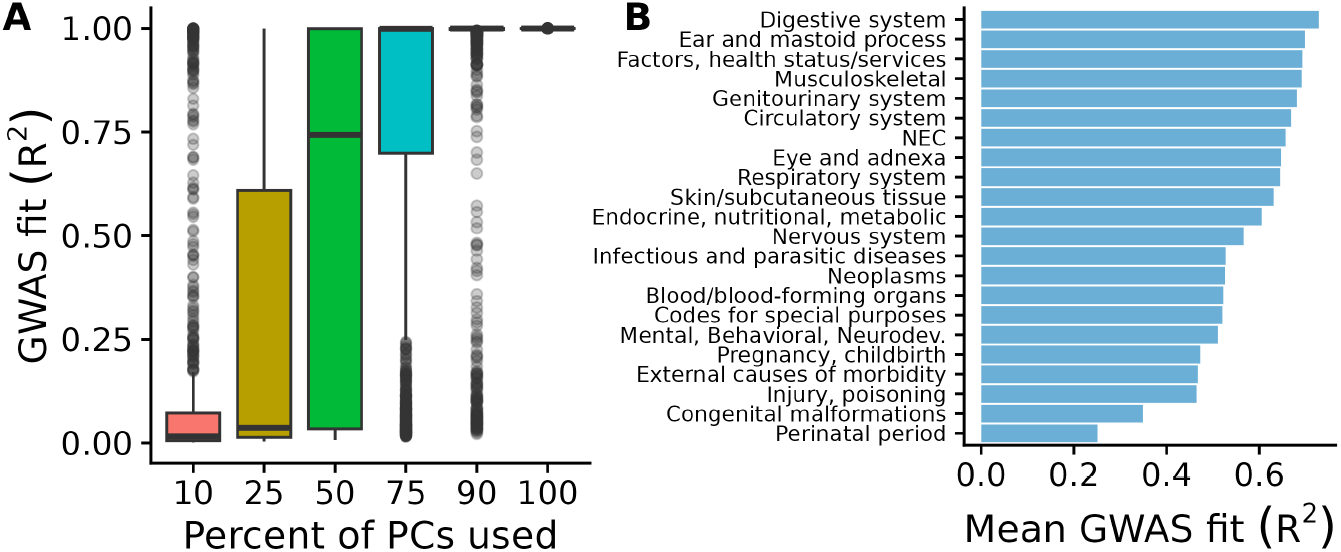
Indirect GWAS performance. **A**. Indirect GWAS performance increases as more PCs are used. Each point represents one phenotype, and the y-axis represents how well the negative log p-values are reconstructed using Indirect GWAS. Some phenotypes can be reconstructed accurately at low latent dimensions (e.g. 10%), while others require very large latent dimensions (*>*90%). **B**. Performance across ICD-10 sections. Performance was defined as the mean across phenotypes and compression fractions within a category. Category names were shortened to allow visualization.

To gain more insight into GWAS performance, we scrutinized the performance of Indirect GWAS across various compression sizes, using phenotype categories provided in the ICD. Specifically, ICD-10 defines 20 sections, which correspond to ranges of codes, such as “A00-B99: Certain infectious and parasitic diseases”. Since the GWAS performance of any individual phenotype only increases as the compression fraction increases (Figure 5A), we computed the performance of each category as simply the mean across compression fractions for each phenotype and across phenotypes within the category. Figure 5B shows the performance for the 20 ICD-10 sections. The best performing section was “K00-K95: Diseases of the Digestive System”, while the worst performing was “P00-P96: Certain Conditions Originating in the Perinatal Period”.

## 3 Discussion

In this work, we introduced Indirect GWAS, a mathematical method for reconstructing GWAS summary statistics for linear phenotype projections using only summary information. Our method aims to improve the speed and flexibility of pan-biobank GWAS by improving computation time and allowing the analysis of custom phenotype definitions, beyond the predefined pan-biobank GWAS phenotypes. To speed pan-biobank GWAS computation, we propose reducing the dimensionality of the phenotypes using a linear method such as PCA, performing GWAS on the lower-dimensional space, and then reconstructing high-dimensional GWAS summary statistics using the linear map from latent to original space. To allow the analysis of custom phenotypes, we propose using linear regression to approximate the chosen phenotype with the phenotypes included in a pan-biobank GWAS as features, applying Indirect GWAS to reconstruct summary statistics, and using the goodness-of-fit for each phenotype as a measure of the fidelity in the summary statistics.

The biggest limitation of Indirect GWAS is its reliance on linear projections. This restricts both the GWAS methods that can be used (e.g. no logistic regression or generalized linear mixed models) as well as the dimensionality reduction methods that can be used (e.g. cannot use deep autoencoders). While linear approximations may or may not perform well for all phenotypes, nonetheless the Indirect GWAS approach can quantify the confidence we have in GWAS summary statistics by measuring the phenotypic goodness-of-fit (Figure 3C). Moreover, as linear and logistic regressions produce summary statistics that match closely for variants with small or moderate effect [10], Indirect GWAS’s limitation here is not critical.

We evaluated the runtime of various methods with and without Indirect GWAS, and using a variety of latent space sizes. Given a sufficient reduction in phenotype dimensionality, Indirect GWAS led to large speed improvements over fully direct methods (Figure 3A). However, different applications could be more or less advantageous to Indirect GWAS. For example, a study of all pairs of phenotypes would be much faster using Indirect GWAS, as phenotype pairs are simple functions of the phenotypes themselves, and pairs could be well-approximated linearly. Conversely, a study of a small number of rare binary phenotypes would be unlikely to benefit much from the Indirect GWAS approach, as these phenotypes are unlikely to be well-approximated linearly with few features.

Another limitation of the current study is that our runtime comparisons are certainly imperfect. Our comparison did not use highly optimized parallel implementations such as SAIGE implemented in Hail [11] or REGENIE in Glow [12]. Our implementation of Indirect GWAS could also be improved, and the majority of its runtime is spent reading and writing GWAS summary statistics from disk. Overall, we report timings to illustrate scaling behavior and to illustrate the advantages of using Indirect GWAS, not to provide an exact reference for runtime expectations.

While Indirect GWAS can provide major reductions in pan-biobank GWAS runtime, these reductions come at the cost of reduced summary statistic fidelity. Nonetheless, we observed that highly-significant associations (low p-values) were generally well-approximated, even at small latent dimensions (Figure 3 and Table 1). As highly-significant variants are generally most interesting for further analysis, this suggests that Indirect GWAS’s overall lower precision is not critical.

Finally, while Indirect GWAS enables computing GWAS summary statistics without access to individual-level data, it requires pre-specified coefficients that can be challenging to pick without individual-level data. Future work could adopt privacy-preserving methods for determining these coefficients using real data (e.g. [13]).

## 4 Methods

GWAS summary statistics primarily include the following pieces of information for each genetic variant tested: coefficient estimate, coefficient standard error, p-value, and sample size. We take the indirect sample size to be the minimum of the sample sizes available for each feature trait, as this is the number of samples that would be available in the equivalent direct approach. We will focus on the coefficient estimate and its standard error, as the p-value can be computed from these statistics.

Consider a regression involving a single genetic variant, a single phenotype, and *N* samples. Let **y** be the *N* -vector of phenotype values, **g** be the *N*-vector of genotype values (coded in any way), and **Z** be the *N × C* matrix of covariates that includes an intercept and is assumed to be full-rank.

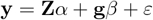

Above, *α* is the *C*-vector of covariate fixed effects, *β* is the scalar effect of the genotype on the phenotype, and *ε* is the *N* -vector of errors. Covariate effects can be removed using a residual projection matrix, **P** = **I**_*N*_ *−* **Z**(**Z**^*T*^**Z**)^*−*1^**Z**^*T*^. Removing covariates results in residualized phenotype 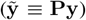, genotype 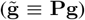, and error 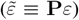 vectors, and it allows writing the regression more compactly.

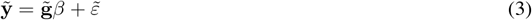

Equation 3 is a univariate least squares regression. Let *d* be the appropriate degrees of freedom for this analysis. Accordingly, estimates of its coefficient 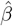 and standard error 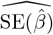 are the following standard results:

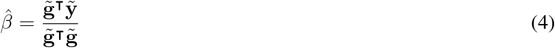

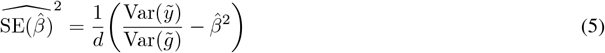

Now, suppose *y* can be written as a linear combination of *m* feature phenotypes. Let **X** be an *N × m* matrix of feature phenotypes and let **p** be an *m*-vector of the projection coefficients.

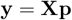

Suppose that for every phenotype in **X**, the same genotype-phenotype regression has been performed. Let 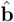 and 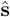 be *m*-vectors holding coefficient estimates and standard errors, respectively, from these regressions (e.g.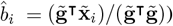 . To estimate the standard error for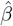, we first need an estimate of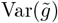, which we obtain by solving equation 5 using 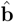 and 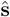

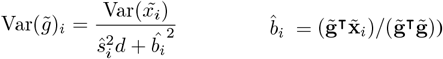

This equation for the genotypic partial variance is written in terms of variables that can be obtained from a GWAS study—the phenotypic partial variance, the estimated coefficient, the estimated coefficient standard error, and the number of degrees-of-freedom (*d* = *N − C −* 1). Note that every input phenotype *i* will result in a different estimate for this value. For Indirect GWAS, we use the mean of these values across phenotypes as the genotype partial variance 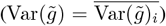 . Unless cohorts are different for the different input GWAS, this value will be the same for every phenotype. The final term that remains to be estimated is 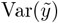, which can be computed using the partial covariance matrix of the feature traits, **C**.

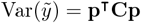

Finally, we can write the coefficient and standard error estimates for *y* as follows:

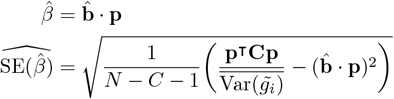

### 4.1 Implementation

Indirect GWAS is mathematically straightforward but challenging to implement efficiently. The method requires potentially thousands of input files, including GWAS summary statistics for every input phenotype, a phenotypic covariance matrix, and a phenotype projection matrix. As the total size of these files can be very large, we provide a high-performance implementation that uses multithreading and chunked processing to provide results efficiently. Our implementation is easily installable, written in Rust, and freely available on GitHub [14].

## 5 Supplementary tables

**Table S1:**
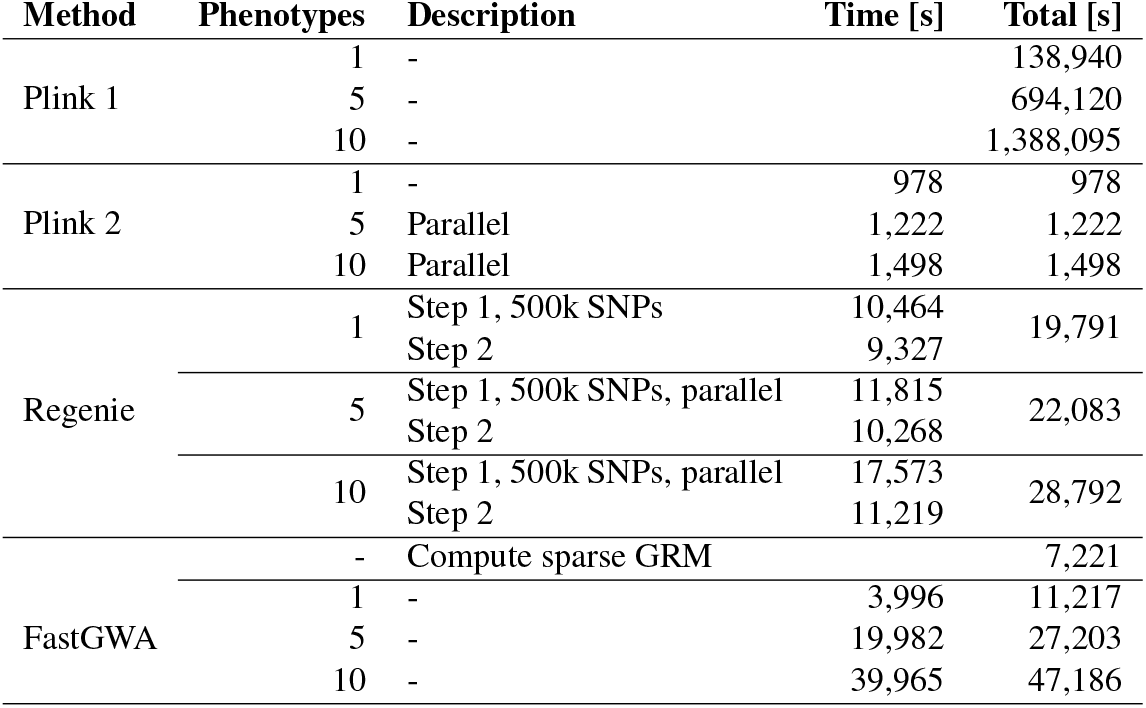
Measured runtime for direct GWAS approaches/steps. Each GWAS involved 342,350 samples, and, unless otherwise noted, 1,166,145 imputed HapMap3 variants. Parallel indicates that multiple phenotypes are run at once, an optimization that improves scaling behavior as more phenotypes are added. Per the recommendations of the Regenie developers, we performed step 1 using a subset of 500,000 variants.

| Method | Phenotypes | Description | Time [s] | Total [s] |
| --- | --- | --- | --- | --- |
| Plink 1 | 1 | - |  | 138,940 |
|  | 5 | - |  | 694,120 |
|  | 10 | - |  | 1,388,095 |
| Plink 2 | 1 | - | 978 | 978 |
|  | 5 | Parallel | 1,222 | 1,222 |
|  | 10 | Parallel | 1,498 | 1,498 |
| Regenie | 1 | Step 1, 500k SNPs | 10,464 | 19,791 |
|  |  | Step 2 | 9,327 |  |
|  | 5 | Step 1, 500k SNPs, parallel | 11,815 | 22,083 |
|  |  | Step 2 | 10,268 |  |
|  | 10 | Step 1, 500k SNPs, parallel | 17,573 | 28,792 |
|  |  | Step 2 | 11,219 |  |
| FastGWA | - | Compute sparse GRM |  | 7,221 |
|  | 1 | - | 3,996 | 11,217 |
|  | 5 | - | 19,982 | 27,203 |
|  | 10 | - | 39,965 | 47,186 |

**Table S2:**
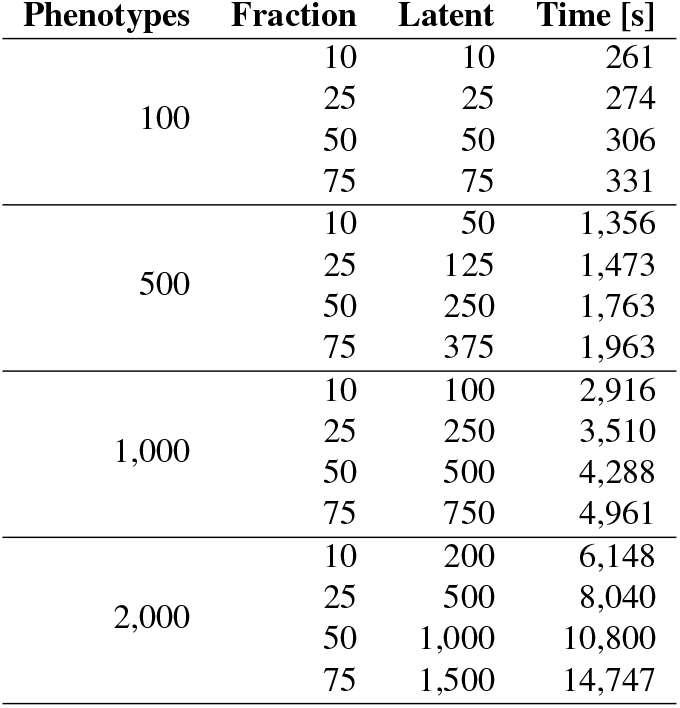
Indirect GWAS runtime results. Phenotypes indicates the number of output phenotypes written by Indirect GWAS. Fraction indicates the latent space size as a fraction of the original size. Latent indicates the number of latent phenotypes. Time indicates the time required to read all the inputs, compute output summary statistics, and save them to a compressed output file. This analysis used summary statistics for 1,166,145 imputed HapMap3 variants.

## 6 Supplementary methods

### 6.1 UK Biobank data processing

To begin, we selected a cohort using only White British individuals to reduce the effects of population structure on our analysis. Next, we removed individuals whose data were flagged for various reasons as being potentially flawed or erroneous. Specifically, we removed individuals whose genetic sex was mismatched with their self-reported sex, individuals who were outliers for heterozygosity or missingness (defined by the UK Biobank using the outlier detection algorithm, *abberant* [15]), individuals with ten or more third degree relatives in the UK Biobank, individuals with sex chromosome aneuploidy, and we restricted to individuals used in the computation of genetic principal components by the UK Biobank. Finally, we restricted to individuals with both binary and quantitative phenotype data available, as described below.

We gathered phenotypic data for this cohort using both binary and quantitative phenotypes. For binary phenotypes, we considered all International Statistical Classification of Diseases and Related Health Problems, 10th revision (ICD-10) codes. In the UK Biobank, these can be obtained from six different data fields, and we included them all. These fields are hospital inpatient ICD-9 codes, hospital inpatient ICD-10 codes, self-reported non-cancer illness codes, primary cause of death, secondary cause of death, and general practitioner outpatient diagnoses. We used mappings provided by the UK Biobank to convert each coding to ICD-10. Only codes with at least 100 observations were retained.

For quantitative phenotypes, we considered all quantitative data fields and excluded fields that were not relevant to this study. Specifically, we chose fields meeting the following criteria: only field data (exclude all bulk data, e.g. imaging), unisex fields only, only fields with continuous or integer values, only fields with at least 75,000 samples (per the August 2023 UK Biobank data dictionary), no strictly genetic fields (e.g. exclude genetic principal components (PCs), exclude pre-computed polygenic scores), no fields dealing with sample quality control (QC) or calibration (e.g. genotyping batch, Affymetrix QC metrics), no fields dealing with home location (e.g. amount of traffic around home location), and no fields dealing strictly with data collection (e.g. number of samples taken). We manually filtered fields using these criteria.

Many fields have both instance and array indices. Instances correspond to visits to the UK Biobank assessment center. Array indices indicate that multiple measurements or values result from a single investigation instance. We kept array indices as separate fields, and took the per-individual mean across instances for each field. Finally, we transformed all quantitative phenotypes using the inverse rank normal transformation, leading to approximately standard normal distributions. Applying the above QC and filtering procedure resulted in 1378 quantitative phenotypes, 1238 binary phenotypes, and 342,350 samples. To save computation time, we restricted to HapMap3 SNPs, resulting in 1,166,145 SNPs in our final dataset.

### 6.2 Runtime measurements

Our overall approach to runtime measurement was as follows. First, estimate runtime as a function of the number of phenotypes for each direct GWAS method. Second, estimate the runtime for Indirect GWAS alone as a function of the number of features (inputs) and projections (outputs) for 10%, 25%, and 50% compression (features vs projections, e.g. 10% is 500 features, 5000 projections). Third, estimate the runtime of a full Indirect GWAS approach as a function of the number of phenotypes and the compression fraction. Measuring the runtime of large-scale GWAS is not trivial, as a number of factors play a role. While we attempted to make all measurements in as comparable an environment as possible, these runtime measurements are presented with the disclaimer that they are conditional on the hardware and software we currently have available, and should be used to inform more about scaling behavior than absolute runtimes.

We first measured the runtime of the following direct GWAS methods: OLS (in Plink 1 and Plink 2), FastGWA (in GCTA [16]), and Regenie [8]. Because the runtime scaling behavior of each method is approximately linear in the number of phenotypes, we measured GWAS time for only a few sets of phenotypes each (N=1, 5, and 10), then extrapolated to estimate the runtime for much larger sets of phenotypes. For this analysis, the 10 phenotypes included were arbitrary; we used the 10 blood biochemistry measurements whose UK Biobank data fields begin with 307. Each runtime was measured using the real time, as reported by GNU Time. All measurements used a UK Biobank cohort of 342,350 individuals and, unless otherwise specified, 1,166,145 imputed HapMap3 variants.

We estimated the runtime of Indirect GWAS using 10, 100, and 1000 phenotypes, each with 10%, 25%, or 50%, compression. This allowed us to extrapolate its runtime to arbitrary configurations. Finally, we estimated the total computation time of the indirect approach. Given *n* phenotypes, a chosen direct GWAS method (e.g. Regenie), and a chosen compression fraction (e.g. 25%), the Indirect GWAS runtime is the time required to compute PCA for *n* phenotypes, plus the time required to compute direct GWAS on the compressed dimension (e.g. *n/*4), plus the time required to reconstruct summary statistics for all *n* phenotypes using compressed (e.g. *n/*4) features.

